# Accumbens cholinergic interneurons mediate cue-induced nicotine seeking and associated glutamatergic plasticity

**DOI:** 10.1101/2020.05.28.119222

**Authors:** Jonna M. Leyrer-Jackson, Michael Holter, Paula F. Overby, Jason M. Newbern, Michael D. Scofield, M. Foster Olive, Cassandra D. Gipson

## Abstract

Nicotine, the primary addictive substance in tobacco, is widely abused. Relapse to cues associated with nicotine results in increased glutamate release within nucleus accumbens core (NAcore), modifying synaptic plasticity of medium spiny neurons (MSNs) which contributes to reinstatement of nicotine seeking. However, the role of cholinergic interneurons (ChIs) within the NAcore in mediating these neurobehavioral processes in unknown. ChIs represent less than 1% of the accumbens neuronal population yet are activated during drug seeking and reward-predicting events. Thus, we hypothesized that ChIs may play a significant role in mediating glutamatergic plasticity that underlies nicotine seeking behavior. Using chemogenetics transgenic rats that express Cre under the control of the choline acetyltransferase (ChAT) promoter, ChIs were bi-directionally manipulated prior to cue-induced reinstatement. Following nicotine self-administration and extinction training, ChIs were activated or inhibited prior to a cue reinstatement session. Following reinstatement, whole-cell electrophysiology from NAcore MSNs was used to assess changes in plasticity, measured via α-amino-3-hydroxy-5-methyl-4-isoxazolepropionic acid (AMPA) / N-Methyl-D-Aspartate (NMDA) (A/N) ratios. Chemogenetic inhibition of ChIs inhibited cued nicotine seeking and resulted in decreased A/N, whereas activation of ChIs had no effect, demonstrating that ChI inhibition prevents transient synaptic potentiation (t-SP) associated with cue-induced nicotine seeking. To assess potential underlying mechanisms, accumbens α_4_β_2_- and α_7_-containing nicotinic ACh receptors (nAChRs) were pharmacologically inhibited and MSN synaptic morphology was assessed following reinstatement. Inhibition of both nAChR subtypes prevented cue-induced nicotine seeking and t-SP (measured via changes in spine head diameter). Together, these results demonstrate that these neurons mediate cue-induced nicotine reinstatement and underlying synaptic plasticity within the NAcore.

## Introduction

Tobacco use disorder is the leading preventable cause of death within the United States and represents a substantial burden to public health [1]. Self-administration of nicotine produces robust cellular adaptations within brain regions associated with drug reward, including the nucleus accumbens core (NAcore) [2–5]. NAcore glutamatergic signaling is involved in nicotine relapse, where high levels of glutamate release driven by nicotine-paired cues alters the synaptic plasticity in medium spiny neurons (MSNs) [2–4]. Previously, we have shown that cue-induced nicotine reinstatement induces transient synaptic potentiation (t-SP) of MSNs, where a 15 min cue-induced reinstatement session increases MSN α-amino-3-hydroxy-5-methyl-4-isoxazolepropionic acid (AMPA) / N-methyl-D-aspartate (NMDA) (A/N) ratios and spine head diameter (d_h_) in a rapid and transient manner [2]. Further, magnitude of t-SP positively correlates with drug seeking [6]. The neural circuitry and mechanisms driving t-SP, however, are not fully understood.

Nicotine modulates nicotinic acetylcholine receptors (nAChRs), which are expressed on cholinergic interneurons (ChIs) in the NAcore. ChIs account for less than 1% of the cell population, they have the ability to exert powerful modulatory control over accumbens circuitry [7, 8] and may play a role in t-SP. ChIs provide most of the intrinsic cholinergic innervation of the NAcore and are widely distributed throughout the striatum [9]. Additionally, ChIs provide cholinergic modulation of striatal dopaminergic transmission and are known to co-release ACh and glutamate [10–13], allowing for additional modulation of MSN activity [14, 15]. Importantly, NAcore ChIs are involved in reward-predicting events [16] and extracellular ACh is elevated following drug intake [17–20]. ACh release within the NAcore parallels intake of cocaine and remifentanil self-administration acquistion, and blocking nicotinic acetylcholine receptors (nAChRs) inhibits drug acquisition [17]. Together, these studies indicate that ChI-induced activation of nAChRs plays an important role in driving motivated drug use.

Increased glutamate release from prelimbic afferents targeting the NAcore increases cue-induced drug seeking [21] by modulating MSN synaptic physiology [2, 22, 23]. Given that NAcore ChIs are involved in drug-motivated behavior, we hypothesized that this small population of cells mediates glutamatergic plasticity and nicotine-seeking behavior. Using choline acetyltransferase cre-recombinase transgenic rats (ChAT::cre), ChI activity was bi-directionally manipulated via viral administration of cre-dependent Designer Receptors Exclusively Activated by Designer Drugs (DREADDS) prior to cue-induced reinstatement. Whole-cell electrophysiological recordings were then conducted from NAcore MSNs to determine if modulation of ChI activity mediates t-SP measured via A/N currents. Due to the known role of nAChRs in nicotine seeking [5, 24–28], we assessed their role in t-SP. Specifically, we determined if intra-NAcore antagonism of two nAChR subtypes linked to nicotine related behaviors, α_7_- and α_4_β_2_ [5, 18, 27, 29], reduced cue-induced nicotine seeking and t-SP measured via changes in d_h_.

## Materials and Methods

### Subjects

Sixty- to ninety-day old adult Long Evans male (N=17) and female (N=20) rats used for chemogenetic ChI experiments and twelve outbred male Sprague-Dawley rats used for nAChR pharmacological experiments were housed in a temperature and humidity-controlled animal facility on a 12-hour reverse light cycle and had ad libitum access to food and water prior to experimentation. All procedures were approved by the Institutional Animal Care and Use Committee (IACUC) of Arizona State University (ChI experiments) or the Medical University of South Carolina (MUSC; nAChR pharmacological experiments). All animals used for chemogenetic ChI manipulation were bred in-house and confirmed ChAT::cre positive through genotyping described in supplemental methods. Breeder ChAT::cre positive males (Long Evans-Tg(ChAT-cre5)5.1 Deis) were purchased from Rat Resource and Research Center (RRRC, RRC#658, Columbia, MO) and bred in-house with Long Evans wildtype females purchased from Envigo (Indianapolis, IN). All Sprague-Dawley rats (250-300g) were purchased from Charles River (Wilmington, MA). Data in Figures 1-3 and S1-S4 were generated at ASU. Data in Figure 4 were generated at MUSC.

**Figure 1:**
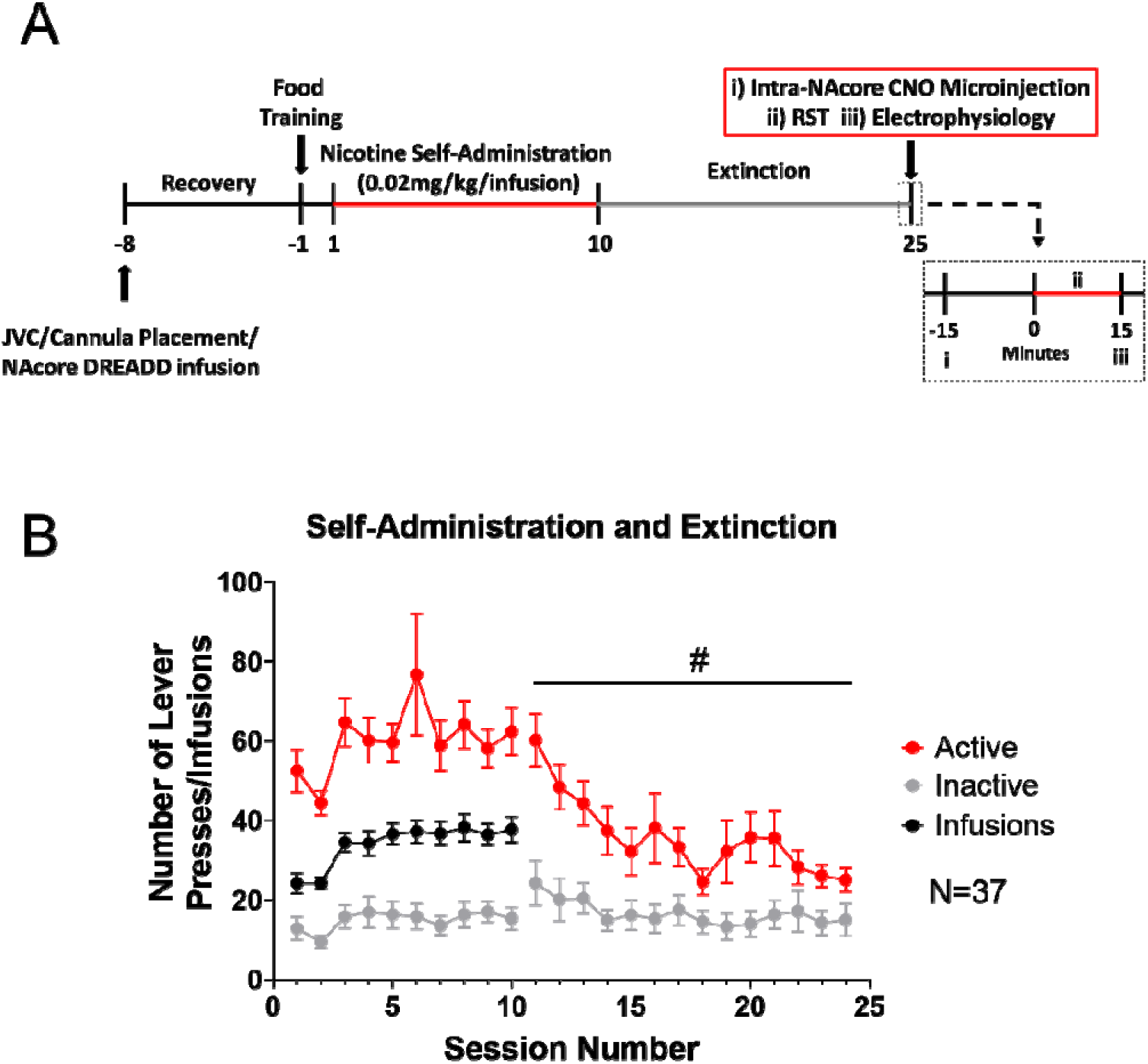
Nicotine self-administration and extinction. (A) A timeline of experimental procedures. JVC = Jugular Vein Catheterization; RST = Reinstatement (B) Rats acquired nicotine self-administration, distinguishing between active (red) and inactive (gray) levers to receive intravenous infusions of nicotine (black). Active lever pressing was significantly reduced across extinction sessions. #*p*<0.05 represents a significant main effect of session on active lever pressing.

All ChAT::cre self-administration data presented here is included within a larger data set exploring the role of sex and strain on nicotine self-administration in our recently published study [30]. However, while these animals were included within the Long Evans male and female groups, the study did not focus on the subset of animals as reported here, but rather included in a larger dataset focusing on self-administration parameters across strain and sex.

### Surgical Procedures

All rats were anesthetized using ketamine hydrochloride (80-100 mg/kg, i.m.) and xylazine (8 mg/kg, i.m.) and underwent surgical implantation of intravenous jugular catheters as well as stereotaxically implanted guide cannulae targeting the NAcore as previously described [30, 31]. Guide cannulae were bilaterally implanted (+1.5 mm anterior/posterior, +/− 2.0 mm medial/lateral, and −5.5 mm dorsal/ventral; [32]). Microinjectors protruded 2 mm past the guide cannulae into the NAcore. Immediately following implantation, viral vectors encoding DREADDs for ChI manipulation were infused into the NAcore of ChAT::cre animals: AAV5-hsyn-DIO-HM4D(G_i_)-mCherry (inhibitory; titer: 1.2×10^13^ vg/mL; N=15 [Addgene, #44362]), AAV5-hsyn-DIO-rM3D(G_s_)-mCherry (excitatory; titer: 1.3×10^13^ vg/mL; N=10 [Addgene, #50485; packaged into an AAV5 vector by Penn Vector Core; Philadelphia, PA, USA]), or AAV5-hSyn-DIO-mCherry (control (i.e. only mCherry-expressing); titer: 1.5×10^13^ vg/mL; N=12 [Addgene, #44362]) at a volume of 0.5 μL per hemisphere. Rats were immediately administered cefazolin (100 mg/kg, i.v.) and heparin (10 U/mL, i.v.) and for 7 consecutive days during the recovery period. Meloxicam (1 mg/kg, s.c.) was given immediately and for the first 3 days of the recovery period. Heparin (10 U/mL, i.v.) was administered daily. Nicotine (0.02 mg/kg/infusion across 5.9 sec) was paired with a compound stimulus (light+tone), and was followed by a 20s time out period. Session duration was 2 hr and a fixed ratio (FR)-1 schedule of reinforcement was utilized. All animals were required to complete a minimum of 10 sessions prior to moving into extinction with the following criteria: ≥10 nicotine infusions obtained and ≥2:1 active/inactive lever press ratio.

During the extinction phase, active lever presses no longer resulted in nicotine infusions or associated cues. A minimum of 14 2-hr extinction sessions was required for each animal. Extinction criteria were set at less than 30 active lever presses on the last day of extinction. For reinstatement sessions, previously paired nicotine cues were presented upon active lever pressing, however no nicotine infusions were delivered. Reinstatement sessions were 15 minutes in duration. Animals were then immediately sacrificed for whole-cell electrophysiological recordings (ChI manipulation experiment) or dendritic spine morphology (nAChR pharmacological experiment). Eight animals were removed from the current study due to not meeting self-administration criteria (N=5) or lack of DREADD expression (N=3). No animals were removed from the nAChR pharmacological experiment, however, two animals were removed from spine analysis due to poor DiI cell impregnation.

### Intra-NAcore Microinjections

For ChI experiments, intra-NAcore microinjections of clozapine-N-oxide (CNO; 0.1 mg/mL dissolved in ACSF) at a volume of 0.5μL were conducted 15 minutes prior to reinstatement testing (Figure 1A). Intra-NAcore microinjections were chosen to avoid potential indirect effects of systemic CNO, since intra-cranial CNO administration does not exhibit DREADD-independent effects in multiple brain regions (for review see [33]). For the nAChR antagonism experiments, intra-NAcore microinjections of the α_4_β_2_ antagonist dihydro-β-erythroidine hydrobromide (DHβE; 140 nmole), the α_7_ antagonist methyllycaconitine (MLA; 16.5 nmole), or vehicle (ACSF) at a volume of 0.5μL were administered 10 min prior the reinstatement session (Figure 4A).

### Whole-cell Electrophysiology

Brains were rapidly removed and submerged in ice-cold carbogen (95% O_2_ / 5% CO_2_) saturated cutting solution (cutting ACSF). Coronal slices containing NAcore were made at 300μm in ice-cold cutting ACSF, and then placed into a continuously oxygenated holding chamber filled with recording ACSF solution for 45 min at 34°C, before allowing to cool to room temperature. All recordings were conducted from NAcore MSNs following visual verification of cannulae placement and initiated 10 minutes after cell membrane rupture. An electrical stimulating electrode was placed in the dorsal region of the NAcore to activate prelimbic excitatory fibers targeting the NAcore. Excitatory post-synaptic currents (EPSCs) composed of both AMPA and NMDA receptor mediated currents were evoked at +40mV. DNQX (20 μM) was then bath applied for 5 minutes and NMDA receptor mediated currents were obtained. AMPA currents were obtained by subtracting the isolated NMDA receptor-mediated current from the whole EPSC. A/N ratios were calculated by measuring the peak amplitude of each current and taking a ratio. For DREADD function validation, see procedures outlined in [34]. All recordings were conducted using the recording software Axograph. Responses were digitized at 10 kHz and saved on a disk using digidata interface (Axon Instruments) and analyzed offline using Axograph.

### Immunohistochemistry

Slices from electrophysiology were post-fixed in 4% paraformaldehyde (PFA) solution and immunohistochemistry was conducted. Slices were probed for ChAT using a mouse monoclonal Anti-ChAT antibody (1:1000; Atlas Antibodies, Bromma, Sweden; AMAB91130) and AlexaFluor 488 conjugated donkey anti-mouse IgG secondary antibody (1:200; Abcam, Cambridge, MA, USA; ab205718).

### Dendritic Spine Morphology

All dendritic spine labeling of NAcore MSNs is described in our previous studies [6, 35] and published protocol [36]. Images were obtained on a Zeiss LSM510 confocal microscope.

### Drugs and Viral Vectors

(−)Nicotine tartrate (MP Biomedicals, Solon, OH, USA) was dissolved in 0.9% sterile saline and adjusted to pH 7.2-7.4 with 1M NaOH. The final stock concentration was 0.2 mg/mL free base, which was adjusted for body weight to achieve an infusion concentration of 0.02 mg/kg/mL. CNO was purchased from Sigma-Aldrich (St. Louis, MO) and diluted in ACSF purchased from Tocris Biotech to 0.1 mg/mL. Heparin, xylazine (100 mg/mL), cefazolin and meloxicam used at 10 U/mL, 8mg/kg/mL, 100mg/kg/mL and 1mg/kg in 0.9% sterile saline, respectively. DHβE and MLA citrate were purchased from Tocris Biomedical Sciences (Minneapolis, MN) and diluted to 140 and 16.5 nmole in ACSF, respectively.

### Data Analysis

Analysis of self-administration, extinction and reinstatement data was performed using two-way, mixed measures ANOVA with DREADD virus as a main factor and session (extinction vs. reinstatement, where applicable). Electrophysiological data were analyzed using a one-way ANOVA, where DREADD virus was considered a factor. Bonferroni-corrected *t* tests post hoc, were conducted where appropriate. The effects of sex were examined in a separate one-way ANOVA to ensure no differences between groups before collapsing for analyses. Analysis of behavioral data only included animals in which met self-administration criteria and expressed DREADDs within the NAcore. Linear regression analyses were used to explore changes in lever pressing across sessions and the relationship between A/N and number of active lever presses. Statistical tests were performed in Graphpad Prism 8.0, and *p*<0.05 was considered statistically significant. Values presented are represented as mean ± standard error of the mean (SEM).

## Results

### Nicotine Self-Administration and Extinction

A two-way ANOVA revealed that active lever pressing remained significantly higher than inactive lever pressing across self-administration sessions (F _9,780_=349.3; *p*<0.05). No interaction between session and lever was observed (*p*>0.05). For extinction training, a two-way ANOVA revealed that active lever pressing significantly decreased across sessions (F_13, 1092_=3.2; *p*<0.05; Figure 1B). Further, a post-hoc comparison revealed that at the beginning of extinction, active lever pressing was significantly higher than inactive lever pressing (t_1092_=5.1; *p*<0.05). However, for the last four sessions, active and inactive lever pressing were not statistically different (*p*>0.05). A linear regression analysis showed a significant difference in the slope of active versus inactive lever pressing, indicating active lever pressing decreased across extinction sessions whereas inactive lever pressing was unchanged (F_1,1116_=9.257; *p*<0.05).

### ChI DREADD validation using immunohistochemistry and electrophysiology

Using immunohistochemistry, all three vectors (two cre-dependent DREADDs and the cre-dependent mCherry-expressing control) were validated to specifically target ChIs. DREADD-labeled neurons (mCherry; figure 2A, left panel) co-expressed ChAT protein (Figure 2A, middle and right panel). Recordings from mCherry tagged DREADD labeled ChIs confirmed functionality of all DREADDs used. Bath application of CNO (10μM) did not alter the resting membrane or spiking in control mCherry-expressing ChIs, induced firing in excitatory G_s_- DREADD-expressing ChIs and inhibited action potential firing in inhibitory G_i-_DREADD-expressing ChIs (Figure 2B).

**Figure 2:**
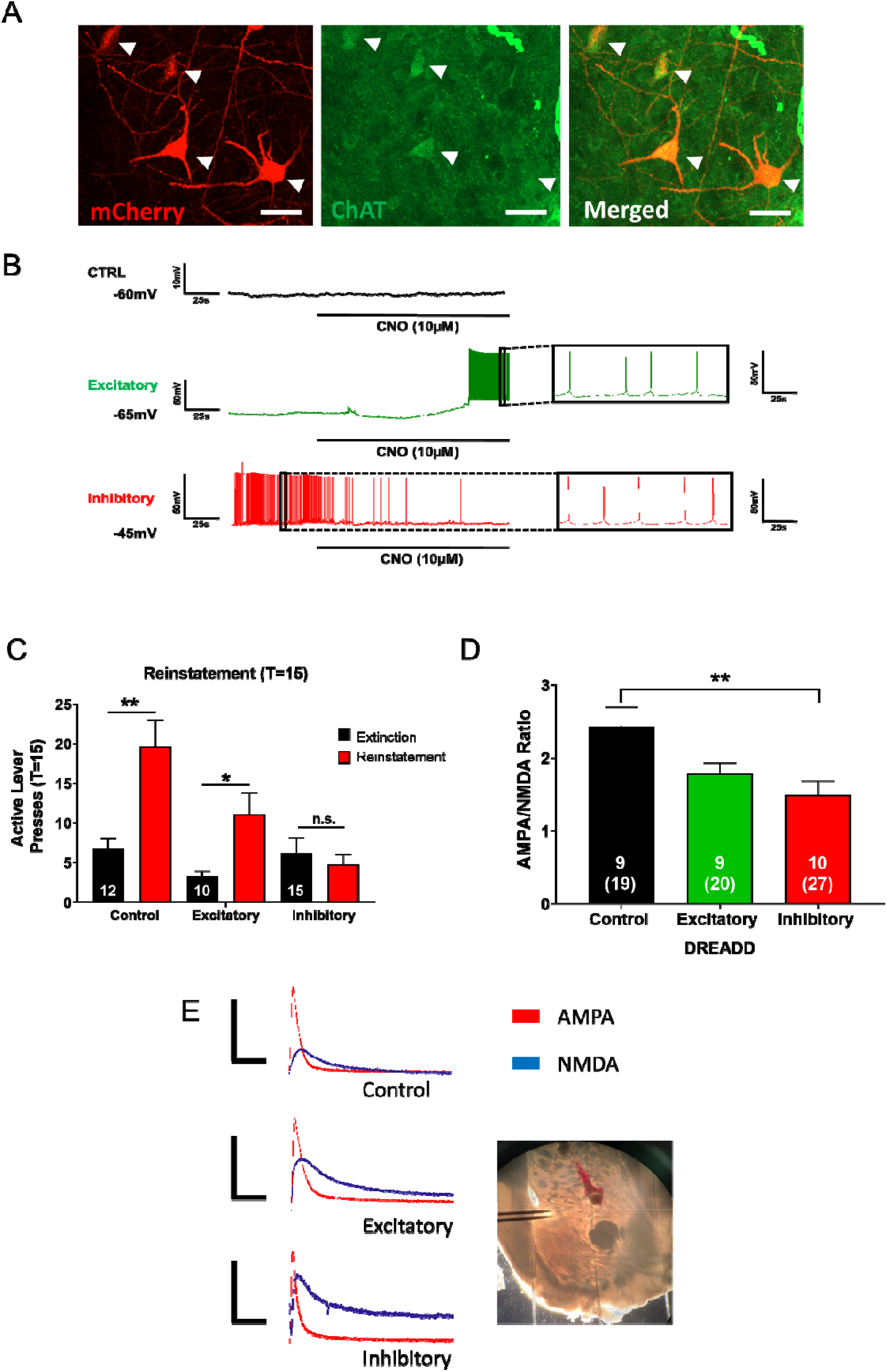
DREADD labeled ChIs co-express choline acetyl-transferase (ChAT). (A) The DREADD constructs used in the current study readily express mCherry in neurons (left panel). ChAT labeled cell bodies within the NAcore (middle panel), which co-expressed with all mCherry labeled neurons (right panel). Arrows depict cell bodies. Scale bar = 30μm. (B) CNO bath application had no effect on control virus expressing ChIs (top; black), promoted firing in the excitatory DREADD-expressing ChI (middle; green); and blunted firing in the inhibitory DREADD-expressing ChI (bottom; red). (C) In control and excitatory DREADD-expressing animals, active lever pressing was significantly increased during cue-induced reinstatement compared to extinction following intra-NAcore CNO treatment. In animals expressing the inhibitory DREADD, CNO inhibition of ChIs prevented cue-induced nicotine reinstatement, where active lever pressing during reinstatement was not different from extinction. * *p*<0.05 vs extinction; ** depicts *p*<0.01 vs. extinction; n.s. = non-significant. Inset numbers represent number of animals. (D) A/N ratio was reduced in animals with ChI inhibition following reinstatement (T(time)=15) relative to control DREADD expressing animals. Representative AMPA and NMDA traces for each DREADD type is shown in panel E. A picture depicting the NAcore and stimulating electrode placement is also shown. Numbers in bars represent animal number and numbers in parentheses represent the total number of cells. Scale bars (x,y) represent = 50ms, 100pA. \*\**p*<0.01.

**Figure 3:**
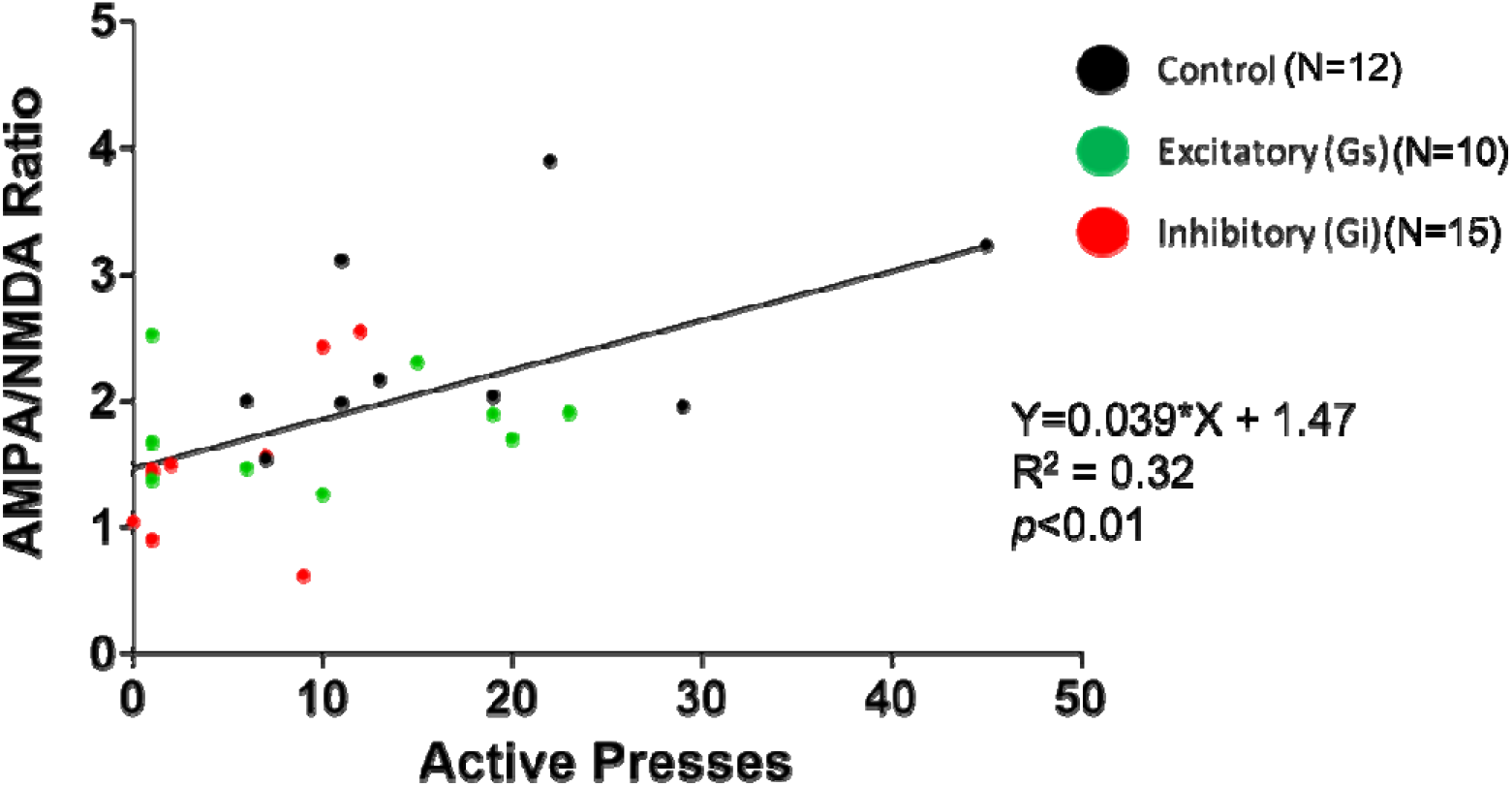
Active lever presses during reinstatement is positively correlated with MSN A/N ratio. The number of active lever presses and A/N ratio for each animal is shown. Group assignment is depicted as follows: control DREADD-expressing rats are shown as black dots, excitatory DREADD-expressing rats are shown as green and inhibitory DREADD-expressing rats are shown as red.

**Figure 4:**
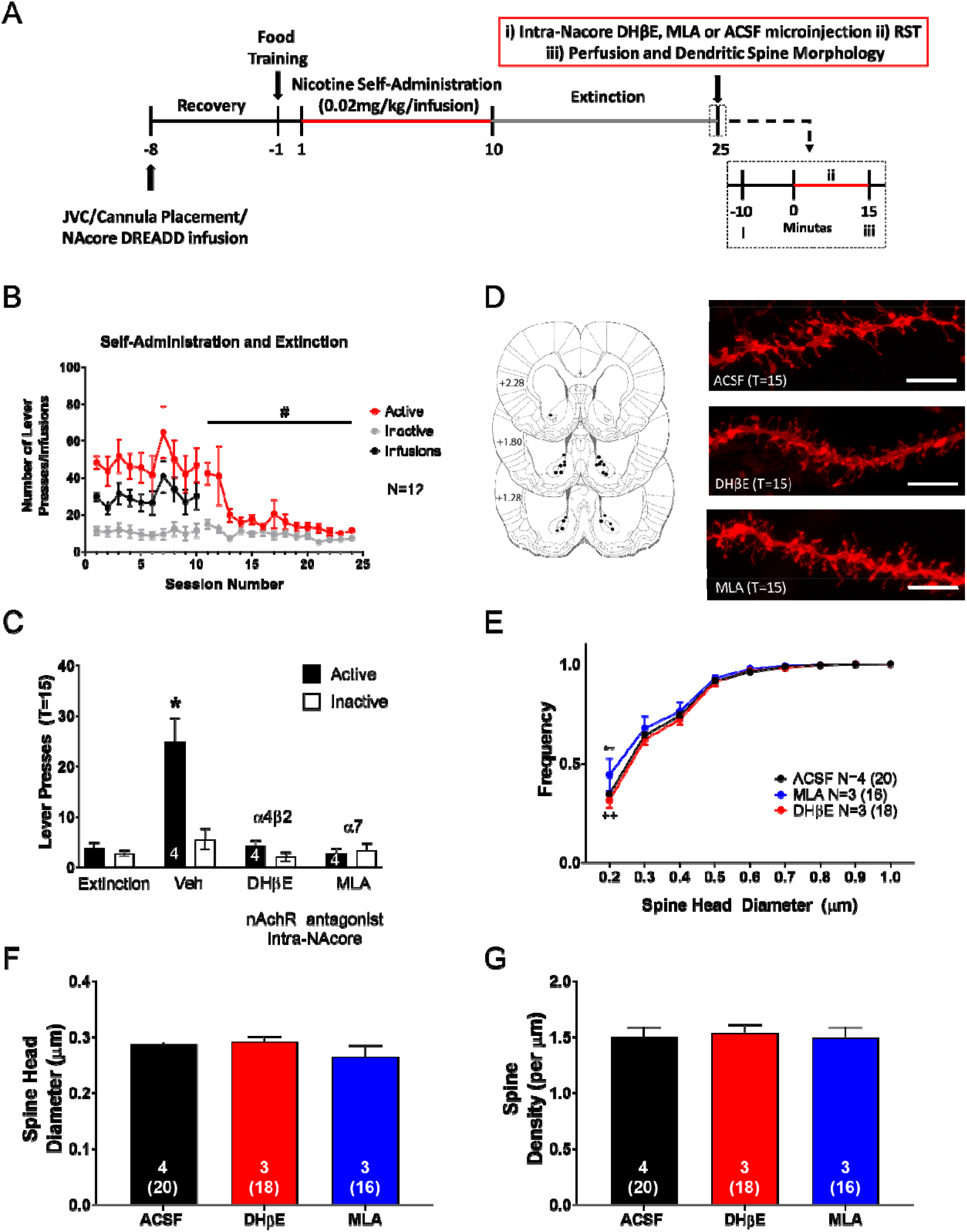
nAChR antagonism prevents cue-induced nicotine seeking and increases the frequency of smaller spine head diameter in MSNs. (A) A timeline of experimental procedures. (B) Rats acquired nicotine self-administration, distinguishing between active (red) and inactive (gray) levers to receive intravenous infusions of nicotine (black). Active lever pressing was significantly reduced across extinction sessions. #p<0.05, significant main effect of session on active lever pressing. (C) ACSF treated animals significantly increased active lever pressing compared to extinction, and antagonism of α_4_β_2_- and α_7_-containing nAChRs, using DHβE and MLA, respectively, prevents cue-induced nicotine seeking. * indicates *p*<0.05 relative to extinction, DHβE and MLA active lever pressing. (D) Histological verification of guide cannula placement according (left) and sample dendritic segments of MSNs from animals treated with ACSF, DHβE and MLA at T=15. Scale bars represent 5 μm. (E) A cumulative frequency distribution of spine head diameter is shown, where MLA significantly increased the number of spines with smaller head diameter relative to ACSF (depicted by ** *p*<0.01) and DHβE (depicted by ++ *p*<0.01). Average spine head diameter and spine head density for groups are shown in (F) and (G), respectively. Numbers in E, F and G represent number of animals and total number of spines in parentheses.

### ChI inhibition prevents cue-induced nicotine reinstatement

A two-way ANOVA with treatment (DREADD type) as between- and session (extinction vs. reinstatement) as within-subject variables revealed a significant main effect of treatment (F_2,70_=8.4; *p*<0.01) and session (F_2,70_=13.2; *p*<0.01). Additionally, an interaction between treatment and session was observed (F_2,70_=6.8; *p*<0.01). Bonferroni post-hoc multiple comparisons revealed that control CNO-treated rats significantly increased active lever pressing during cue-induced reinstatement (T=15) relative to the first 15 min of extinction (t_70_=4.4; *p*<0.01; Figure 2C). Additionally, ChI activation with intra-NAcore administration of CNO did not prevent cue-induced reinstatement in animals expressing the excitatory DREADD, as the number of active lever presses during the cue-induced reinstatement session (T=15) was significantly higher than that observed during the first 15 minutes of extinction (t_70_=2.2; *p*<0.05; Figure 2C). However, inhibition of ChIs with CNO prevented cue-induced nicotine reinstatement in inhibitory DREADD-expressing animals, where the number of active lever presses during reinstatement (T=15) was not different from the first 15 minutes of extinction (*p*>0.05; Figure 2C). No sex differences in active lever pressing within groups were observed (*p*>0.05).

### Inhibition of ChIs prevents t-SP in MSNs

NAcore MSNs recordings were made from acute slices derived from animals expressing either control, inhibitory, and excitatory DREADD constructs immediately following a 15 min nicotine cue-induced reinstatement session. An ANOVA revealed a significant effect of treatment (DREADD type) on A/N (F_2,25_=5.7; *p*<0.01). Post-hoc comparisons revealed that inhibition of ChI activity resulted in a significantly smaller A/N relative to control animals following the 15 min cue-induced reinstatement session (t_25_=3.3; *p*<0.01; Figure 2D). However, no significant differences were observed between rats in which ChIs were activated and control animals (*p*>0.05; Figure 2D) or rats in which ChIs were inhibited (*p*>0.05; Figure 2D). A one-way ANOVA revealed no differences in NMDA decay time between control, excitatory or inhibitory DREADD-expressing animals, as measured by the time to reach 37% of peak amplitude (*p*>0.05; Figure S4C). No sex differences in A/N or NMDA decay within groups were observed (*p*>0.05).

### Reinstatement active lever pressing is positively correlated with MSN A/N ratio

The number of active lever presses and A/N were plotted for each animal. The A/N ratio measured for each cell was averaged across all cells from the same animal (between 1-4 cells/animal) to obtain the A/N for each animal. A linear regression analysis revealed as significant relationship between the number of active lever presses during reinstatement (T=15) and the A/N (F_1,26_=12.4; *p*<0.01; Figure 3). No sex differences in the slope were observed (*p*>0.05).

### Inhibition of α_7_-containing nAChRs prevents cue-induced reinstatement and t-SP in MSNs

A two-way ANOVA revealed that active lever pressing remained significantly higher than inactive lever pressing across self-administration sessions (F _9,220_=144.8; *p*<0.05). However, no interaction between session and lever was observed (*p*>0.05). For extinction training, a two-way ANOVA revealed that active lever pressing significantly decreased across sessions (F_13, 308_=4.7; *p*<0.05; Figure 4B). Further, a post-hoc comparison revealed that at the beginning of extinction, active lever pressing was significantly higher than inactive lever pressing (t_308_=4.8; *p*<0.05). For the last twelve sessions, active and inactive lever pressing were not statistically different (*p*>0.05). A linear regression analysis showed a significant difference in the slope of active versus inactive lever pressing, indicating that active lever pressing decreased across extinction sessions whereas inactive lever pressing was unchanged (F_1,332_=13.4; *p*<0.05). To examine whether blockade of nAChRs inhibited reinstatement, a two-way ANOVA with treatment (ACSF, DHβE or MLA) as between-subject and session (extinction vs. reinstatement) as within-subject variables revealed a significant main effect of treatment (F_2,18_=18.5; *p*<0.01) and session (F_1,18_=12.8; *p*<0.01). Additionally, an interaction between treatment and session was observed (F_2,18_=10.7; *p*<0.01). Bonferroni post-hoc multiple comparisons revealed that following ACSF treated rats significantly increased number of active lever pressing during cue-induced reinstatement (T=15) relative to the first 15 min of extinction (t_18_=5.8; *p*<0.01; Figure 4C). DhβE- and MLA-treated animals did not differ in the number of active lever presses during cue-induced reinstatement (T=15) relative to extinction (*p*>0.05). Further, ACSF-treated animals had significantly higher active lever pressing than animals treated with DHβE (t_18_=6.3; *p*<0.01) and MLA (t_18_=6.7; *p*<0.01; Figure 4C).

Spine morphological measures were analyzed at T=15 following cue-induced reinstatement sessions. Histology and sample dendritic spine segments for each treatment group are shown in Figure 4D (left and right, respectively). A two-way ANOVA with treatment and bin as main factors was used to compare frequency distributions for d_h_. A main effect of treatment (F_2,63_=3.7; *p*<0.05) and bin (F_8,63_=283.9; *p*<0.001) but not interaction between treatment and bin was observed (*p*>0.05). A Bonferroni post-hoc comparison showed a significant increase in the number of spines with smaller spine head diameters following MLA treatment compared to ACSF (t_63_=3.1; *p*<0.01) and DHβE ((t_63_=3.8; *p*<0.01) (Figure 4E). However, no differences in mean d_h_ or spine density were observed between groups (*p*>0.05; Figure 4F and G, respectively).

## Discussion

Here we found that chemogenetic inhibition of NAcore ChIs inhibited cue-induced nicotine seeking behavior. Additionally, inhibition of ChIs reduced A/N relative to control animals receiving the same dose of CNO. Further, the magnitude of reinstated nicotine seeking was positively correlated with the magnitude of t-SP, similar to previous studies [6]. Inhibition of α_4_β_2_- and α_7_-containing nAChRs also prevented cue-induced nicotine reinstatement, however, only inhibition of α_7_-containing nAChRs prevented NAcore t-SP-associated decreases in the frequency of smaller spine heads relative to ACSF-treated animals. Together, these results demonstrate that accumbens ChIs exert significant control over cue-induced nicotine seeking behavior and MSN synaptic physiology within the NAcore, likely through ACh activation of NAcore α_7_-nAChRs located on glutamatergic terminals [37].

### Interactions between ChIs, ACh, nAChRs, and Drug-Motivated Behavior

Within the NAcore, ACh-releasing ChIs exert powerful modulatory control over MSNs and dopaminergic tone [12, 38]. Elevated levels of ACh have been observed in the nucleus accumbens following cocaine [17, 19], remifentanil [17], nicotine [18, 20], and alcohol [39]. Further, elevated levels of ACh in the accumbens parallel the reinforcing effects of both intravenous cocaine and remifentanil [17], providing evidence for ACh mediation of drug-motivated behavior. Interestingly, Ach release in the accumbens is also enhanced during contingent versus non-contingent intravenous cocaine administration, suggesting that ACh release may be differentially mediated by volitional versus passive drug exposure [40]. ChIs are also heavily implicated in stimulus-response associations [41–44], thus it is not surprising that silencing accumbens ChIs prevents cocaine conditioned reward [14]. Witten and colleagues suggest that acute silencing of accumbens ChIs disrupts drug-related learning without affecting conditioning [14]. Thus, while ACh release is primarily driven by activation of ChIs within the accumbens, acute silencing of ChIs in this region can disrupt drug-related learning. In the current study, inhibition of ChIs prevented cue-induced nicotine reinstatement, suggesting that inhibiting ACh release may prevent nicotine seeking in part by blunting the stimulus-response association.

In addition to ChIs, glutamatergic and dopaminergic terminals express nAChRs within the NAcore [9, 11, 45, 46] allowing for extensive mediation of accumbens circuitry. Given the density of nAChRs expressed throughout the NAcore, these receptors have been heavily implicated in the molecular aspects mediating addiction. For example, nAChRs are upregulated within the accumbens following voluntary ethanol consumption [39]. Moreover, nAChR inhibition also reduces cocaine place preference and cocaine sensitivity [47] and nAChR antagonism within the ventral tegmental area attenuates cue-induced cocaine seeking [48]. While the role of nAChRs has been explored across multiple drugs of abuse, their involvement in nicotine use and addiction has been most prominently studied. Nicotine directly activates and leads to prolonged desensitization of nAChRs, which contributes to the reinforcing properties of nicotine [18, 26, 28, 29, 49]. Multiple studies have shown that antagonism of nAChRs reduces active lever pressing in animals self-administering intravenous nicotine [50–52] and antagonism with DHβE blocks the rewarding properties of VTA-administered nicotine [53]. In line with these findings, we have shown that intra-NAcore antagonism of both α_4_β_2_ and α7 containing nAChRs, with DHβE and MLA, respectively, blocked cue-induced nicotine reinstatement (for issues regarding specificity of these compounds see [54–57]). These results are consistent with others demonstrating that inhibition of nAChRs using systemic administration of mecamylamine reduces active lever pressing in animals self-administering nicotine [24, 50, 58]. Together, these results support the hypothesis that the NAcore cholinergic system plays a critical role in nicotine addiction.

### ChI Mediation of t-SP and Nicotine Seeking

NAcore t-SP is characterized by rapid changes in MSN physiological and morphological properties. Specifically, rapid increases in MSN A/N and d_h_ are indicative of t-SP and appear 15 min following a cue-induced nicotine reinstatement session [2]. Here we show that ChI inhibition and antagonism of α_7_-containing nAChRs prevents cue-induced reinstatement and t-SP by suppressing MSN A/N and the decrease in number of smaller spines, respectively. These data suggest that ACh release from ChIs mediates t-SP, likely through mediation of glutamate release via terminally-expressed α_7_-containing nAChR activation [28, 46]. Based on these findings, we hypothesize that ChIs are activated by glutamate release during drug seeking behavior, which may enhance drug seeking behavior through an ACh-induced feedforward mechanism. Specifically, we hypothesize that ChIs release ACh, which then activates glutamate terminal-expressed α_7_-containing nAChRs. Given that these receptors enhance glutamate release [27, 46], we hypothesize that this exacerbates glutamate overflow, t-SP, and nicotine seeking behavior. Within this circuit, our current results likely reflect a reduction in ACh release due to chemogenetic ChI inhibition, leading to a blockade of glutamate release-stimulating α_7_-containing nAChR activation. Although not directly studied here, we hypothesize that when activated by glutamate, ChIs activate presynaptic α_7_-containing nAChRs via ACh to further promote glutamate release. In support, α_7_-containing nAChRs are extensively located on glutamatergic terminals and mediates glutamate release [37, 59]. α_4_β_2_-containing nAChRs are located on dopaminergic terminals and mediate dopamine release within the accumbens [60, 61]. Interestingly, α_4_β_2_-containing nAChRs gate activity-dependent dopamine release and mediate local accumbens circuitry to a greater extent in females, likely driving enhanced motivation for drug seeking compared to males [62]. In the current nAChR study, sex differences were not explored thus it is possible that females exhibit more pronounced changes in MSN dendritic morphology than males. Further, inhibition of α_4_β_2_-containing nAChRs could prevent nicotine reinstatement associated t-SP in females. However, we hypothesize that inhibition of NAcore α_4_β_2_-containing nAChRs with DhβE in the current study with males inhibited dopamine release and prevented cue-induced nicotine seeking, but did not prevent t-SP due to its inability to prevent glutamatergic overflow onto MSNs.

Interestingly, chemogenetic activation of ChIs did not promote reinstatement or enhance A/N beyond control animals. This may be due to ChI autoregulation, where ChI overexcitation and enhanced ACh release induced by CNO may lead to activation of muscarinic M4 autoreceptors located on ChIs. Activation of these receptors induces membrane hyperpolarization and inhibition of ChI calcium channels [9], allowing for self-regulation and a blunting of further ACh release. Due to overexcitation of ChI activity, autoregulatory mechanisms may have prevented ChIs from promoting glutamate release and t-SP. ChIs are the only neuronal cell type within the striatum that express I_h_ (hyperpolarization-activated cation currents, [63, 64]), and contribute substantially to neuronal excitability [65, 66]. Thus, the G_s_ DREADD vector used in the current study may more closely mimic physiological activity by modulating I_h_ compared to G_q_ DREADD vectors. However, endogenous ChI activity not driven by DREADD excitation and subsequent autoregulation may allow for the feedforward glutamate-releasing mechanism to occur, leading to enhanced nicotine seeking behavior and t-SP.

These studies uncover key components of a circuit not previously known to contribute to NAcore t-SP or motivated nicotine seeking behavior. As mentioned above, one limitation of these studies is the lack of direct measurement of ACh release during reinstatement. As well, exploration of ChI autoregulatory mechanisms in relation to t-SP is warranted. Further, future studies should characterize the role of ChI signaling specifically on D1- and D2-expressing MSNs, as t-SP appears to be preferential to the D1 subtype [67]. Although not measured here, these are important next steps to fully characterize the neural circuitry involved in nicotine seeking behavior.

In conclusion, we report that ChI inhibition prevents cue-induced nicotine reinstatement and blunts MSN A/N. Additionally, inhibition of both α_4_β_2_- and α_7_-containing NAcore nAChRs blocks cue-induced nicotine reinstatement, but only inhibition of α_7_-containing nAChRs prevents morphological t-SP. Taken together, these results suggest that the cholinergic system heavily modulates nicotine seeking behavior and associated glutamate plasticity. Thus, ChIs may be an essential pillar of t-SP that has been previously overlooked. Taken together, the current findings illustrate an additional component of the highly complex neurophysiological underpinnings of nicotine relapse, where ChIs mediate control of t-SP and thus play a pertinent role in transitioning nicotine craving to seeking.

## Acknowledgements

The authors would like to acknowledge Neringa Stankeviciute, Nic Allen, Hanaa Ulangkaya, Ngoc Van Do, Jose Piña, Meghan Brickner and Vincent Carfagno for their technical assistance. As well, we would like to thank Dr. Peter Kalivas for use of equipment for the study conducted at MUSC.

## Author Contributions

JMLJ and CDG were responsible for study concept and design. JMLJ, CDG, MH, PO, MS and JN contributed to data collection and data analysis. JMLJ, FO and CDG contributed to interpretation of findings. JMLJ drafted the manuscript. All authors provided critical revision of the manuscript for content and approved the final version for submission and publication.

